# Insights into Immune Gene Prediction and Function Through the Evolutionary History of *ADF* Gene Family

**DOI:** 10.1101/2024.05.31.596878

**Authors:** Huan Chen, Brad Day

## Abstract

- ACTIN DEPOLYMERIZING FACTORS (ADFs) are key regulators of actin cytoskeletal dynamics and plant immunity.
- We predicted the potential immune-associated function of 38 genes from *Arabidopsis* using gene expression values from 24,123 RNA-Seq datasets and 34 single-cell datasets through machine learning algorithms.
- The evolutionary relationships of *ADF* family members from 38 eukaryotic species were evaluated, including an assessment of the sub-function(s) of these members.
- Our results show that the *ADF* clade in plant and other kingdoms are separated, with *ADF3, 5, 7, 9,* and *10* possessing collinear relationships within species, and *ADF 2,3,4,6,7, and 10* possessing evolved, new, sub-functions related to response to Fe, copper-deficiency, and ABA signaling in *Arabidopsis*. Expanded, multiple, roles for *ADF1,4,* and *6* were also identified.
- This study not only provides an analysis of the expanded role for the ADF family of genes/proteins, but also provides insight into, and a framework for, the identification and study of the evolutionary history of genes having putative roles in immune signaling.

## Introduction

*Arabidopsis* is a model species for the study of crop plants, with inherent advantages being a small genome size and a short generation (Arabidopsis Genome, 2000). With 25,498 predicted protein-coding genes, 69% of these have predicted functions based on sequence comparisons and similarities to known proteins. However, approximately only 9% of the genes have been studied or characterized through experimental methods. The remaining 30% remain without any predicted function (Wigge and Weigel, 2001). Although the function, or functions, of genes described to date have been evaluated in large part through advanced sequencing technologies, such as the case of the immune genes ACTIN DEPOLYMERIZING FACTORS (ADFs) (Andrianantoandro and Pollard, 2006, Pavlov et al., 2007), experimental methods have proven time-consuming. These limitations lend themselves to the need to develop new methods to develop testable hypotheses based on more rapid, machine-learning-based, approaches.

The actin cytoskeleton plays a central role in all eukaryotes, regulating various cellular functions in multiple cellular processes, including plant growth, response to environmental changes, cell shape, division and differentiation, facilitating cytoplasmic streaming, organelle movement, and regulation of polarity and stomatal movement (Henty-Ridilla et al., 2013, Staiger and Blanchoin, 2006, Pollard and Cooper, 2009, Hussey et al., 2006, Szymanski and Cosgrove, 2009, Day et al., 2011, Staiger, 2000, Zhao et al., 2011). The actin cytoskeleton is not only highly organized, but also highly dynamic within cells. Indeed, rapid reorganization and turnover is achieved through tightly regulated transitions between globular (G) and filamentous (F) forms of actin, which in total is regulated by 50-200 actin-binding proteins (ABPs), depending on species (Staiger and Blanchoin, 2006). Among the numerous actin-binding proteins that regulate actin organization and dynamics, and thus various biological functions (Henty-Ridilla et al., 2013, Huang et al., 2015, Higaki et al., 2007, Sun et al., 2013, van Gisbergen and Bezanilla, 2013), the ACTIN DEPOLYMERIZING FACTORS (ADFs) family are an ancient protein family that is conserved across all eukaryotes. ADF proteins play a crucial role as actin-binding proteins (ABPs) in eukaryotes, irrespective of their capability to depolymerize F-actin. Indeed, they produce filament ends conducive to new filament initiation, thereby maintaining the balance between F-actin and G-actin. Additionally, ADF enhances the dynamism of actin, as noted by Andrianantoandro and Pollard (2006) and Pavlov et al. (2007).

Beyond their major function as a depolymerization factor of actin microfilaments, plant *ADFs* are involved in various biological process, including hypocotyl elongation, innate immune signaling, response to infection, polar cell growth, stomatal movement, and response to abiotic stress (Dong et al., 2001, Clement et al., 2009, Tian et al., 2009, Zheng et al., 2013, Henty-Ridilla et al., 2014, Inada et al., 2016, Zhao et al., 2016, Zhu et al., 2017, Qian et al., 2019). With such a breadth of roles in numerous plant processes, it is not surprising that the plant *ADF* family is both large, as well has expanded to include multiple functions per member. For example, the *Arabidopsis ADF* family consists of 11 functional genes, which can be divided into four subclasses (subclasses I– IV) (MacRobbie and Kurup, 2007). Within this, further differentiation of expression and function is predicted to exist based on tissue distinctions. For example, *ADF* subclass I in *Arabidopsis* includes *ADF1*, *ADF2*, *ADF3*, and *ADF4*, all of which are expressed throughout the plant at relatively high levels. However, *ADF7* and *ADF10*, both of which are members of subclass II, are only expressed in flowers; specifically, in pollen. Conversely, *ADF8* and *ADF11*, both members of subclass IIb, are only expressed in root epidermal cells, while members of subclass III and subclass IV are expressed in a wide variety of tissues; the expression of *ADF5* and *ADF9* are not found in the root apical meristem (Ruzicka et al., 2007). Results from Dong et al. (2001) have also confirmed the distinct expression pattern of *ADF1*, *ADF5*, and *ADF9*, again with a restricted pattern of expression for *ADF5*.

In addition to the distinct tissue-specific patterns of expression for members of the *Arabidopsis ADF* family, physiological – and functional – role differences have also been reported. For example, and as a demonstration of a role for actin abiotic signaling responses, *ADF1* has been shown to play an important role in plant thermal adaptation by blocking the high-temperature-induced stability of actin filaments (Wang et al., 2023). As a demonstration of the role of actin in plant biotic signaling, there are numerous examples, including one of the earliest which showed that transient expression of *ADF1*, *ADF5*, *ADF6*, *ADF7*, and *ADF10* inhibit host resistance (barley) against powdery mildew (Miklis et al., 2007). More recent studies into the link between actin and plant-biotic interactions revealed a role for *ADF4* as a regulator of gene-for-gene resistance activation and MAPK-signaling via the coordinated regulation of actin cytoskeletal dynamic and *R*-gene-mediated immune signaling (Porter et al., 2012, Peng and Huang, 2006).

In recent years, a key focus on how actin converges on immunity has emerged – specifically, the identification and characterization of shared processes that are influenced by the activity of ADFs and the plant response to infection. At a fundamental level, one of the key first insights into this process came from work by Henty and colleagues which showed that *ADF4* contributes to the stochastic dynamic turnover of actin filaments in the cortical array of epidermal cells (Henty et al., 2011). This work established a foundation for quantitatively evaluating the genetic and physiological interactions that underpin changes in the actin cytoskeleton. From this, studies emerged that further delineated the plant response to environmental, developmental, and physiological stimuli. For example, in the area of abiotic signaling, it has been shown that *ADF5* plays a role in drought stress through regulating stomatal closure and acting as a downstream target gene of CBFs in response to low-temperature stress (Qian et al., 2019, Zhang et al., 2021). Similarly, several recent studies have shown that actin participates in key processes that further converge on the regulation of actin dynamics and response to the environment; these include a role for *ADF7* in pollen tube growth, the inhibition of VLN1, and the activation of signaling in response to osmotic stress (Zheng et al., 2013, Bi et al., 2022). As a demonstration of actin’s role in development, numerous reports demonstrate a role for *ADF9* in flowering (Tholl et al., 2011, Burgos-Rivera et al., 2008), a role for *ADF10* the regulation of vesicle trafficking and pollen tube growth (Jiang et al., 2017). Importantly, while both *ADF7* and *ADF10* are expressed in pollen, and are members of subclass II, it is interesting to hypothesize that they exhibit distinct roles based on differences in expression and localization (Bou Daher et al., 2011, Daher and Geitmann, 2012). Conversely, subclass I *ADFs* play a role in the regulation of nuclear organization and gene expression and is a novel regulator of endoreplication (Matsumoto et al., 2023, Inada et al., 2021).

Recent experimental evidence has contributed to the study of ADF thanks in large part to the coupling of next-generation sequencing technologies, the development of quantitative methods in cell biology, and a robust collection of protein properties datasets. Herein, we predicted the potential immune-associated gene functions from *Arabidopsis* using the *ADF* gene family as a study case; this is based on the duality of their role(s) in structural, signaling, and stress response processes in eukaryotes. Through collection and analysis of the expression profiles of 24,123 RNA-Seq samples and 34 single-cell datasets information from *Arabidopsis*, we analyzed the expression profiles of 11 functional *ADF* genes from different tissues, cell types, as well as in response to different stresses. This approach uncovered the potential immune-associated function gene prediction, the evolutionary history of *ADF* gene family and the sub-function evolution of *ADF*. The results of this study will promote our understanding for the *ADF* gene family and provide the basis for future study of ADFs in plants.

## Materials and Methods

### Prediction of potential immune-associated function genes

A total of 24,123 published RNA-Seq sample of *Arabidopsis thaliana* (L.) were collected from GEO (Gene Expression Omnibus), Trimmomatic (Bolger et al., 2014) were used to remove the adopter and quality control with LEADING:20 TRAILING:20 SLIDINGWINDOW:4:20 MINLEN:20 on the fastq data, salmon (Patro et al., 2017) were used to call Transcripts Per Million (TPM) value from RNA-Seq data. PCA (Principal Component Analysis) was used to convert high-dimensional data into low-dimensional and keep the main information from the gene expression TPM value of 24,123 RNA-Seq samples and total 16 PCs were generated. The information of gene cluster, cell type, tissue were extracted from arabidopsis_thaliana.marker_fd.csv on scPlantDB (He et al., 2024). The missing value in the sing-cell file were filled by using the next value in a backward direction. Genes with information from sing-cell dataset were kept for the machine learning model training.

The correlation was calculated between all the features, and features with correlation number >0.5 were removed prior to training the models (Fig. S1). A total of 3,107 genes from relative biological processes from previous research were used as positive samples, and the sample number of no immune genes was used as a negative sample (Li et al., 2020). Four different algorithms, k-nearest neighbors (KNN) (Dasarathy, 1991), decision tree (Breiman, 1984), random forest (Ho, 1995) and neural networks (NNs) (Hardesty, 2017) were used to train the models. The final predicted immune genes are the common gene from these four different models. All code is pending under GitHub: https://github.com/chenh9313/Sup_ADF.git.

### Species tree construction

Species tree construction among 38 eukaryotic species with 1 bacteria species as outgroup. The protein sequence data of selected species, which presents each phylum of eukaryotes, were download from Ensembl and Ensembl Plants database (https://useast.ensembl.org/index.html; http://plants.ensembl.org/index.html). The species were built using all orthogroups by OrthoFinder (Emms and Kelly, 2015, Emms and Kelly, 2019). The same method as above was used to build the species tree in eight representative species. The representative species were selected based on the genome having been fully annotated and representing significant temporal spacing from common ancestry.

### Phylogenetic tree construction of *ADF* gene family

The phylogenetic tree was performed using the maximum likelihood (ML) method in RAxML (Randomized Axelerated Maximum Likelihood) (Stamatakis, 2014), with 1,000 bootstrap replicates. Sequences of ADF proteins from selected species were identified based on the orthogroups genes results using OrthoFinder (Emms and Kelly, 2015, Emms and Kelly, 2019). Only the longest protein sequence from the longest isoform were used to build the phylogenetic tree. The *ADF* gene from bacterium was found by using *ADFs* genes from *Arabidopsis* to do the Blastp (Altschul et al., 1997) with E value < 0.05. The above methods were also used in the phylogenetic tree construction of ADFs in eight representative modern species and ADFs in *Arabidopsis*. ML phylogenies tree was inferred and visualized by FigTree v1.4.4 (http://tree.bio.ed.ac.uk/software/figtree/).

### Motif analysis of *ADF* gene family in eight representative species

Conserved motifs of ADFs among eight species were predicted using the Multiple Em for Motif Elicitation (MEME) (https://meme-suite.org/meme/) with the following parameters: classic mode, the number of motifs eques 10, the size of motif between 6 wide and 50 wide (inclusive), A 0-order background model generated from the supplied sequences and zero or one occurrence (of a contributing motif site) per sequence.

### Cis-acting elements of the *ADF* gene family

The discovery of cis-acting elements was based on the definition of the promoter region as being a short region of DNA (ca. 100-1,000 bp) typically located directly upstream, or at the 5′ end, of the transcription initiation site, (Le et al., 2019, Lin et al., 2019). Genes contain common motifs in their promoter regions have similar expression patterns in low-oxygen response in *Arabidopsis* root (Vilo et al., 2000, Klok et al., 2002). Thus, the 1,000 bp upstream of the transcription start site of all *ADF* transcripts was extracted as promoters to predict cis-acting elements using PlantCARE (Lescot et al., 2002).

### Chromosomal localization, gene duplication, and calculating Ka/Ks values of ADFs

All the *ADF* genes were mapped to chromosomes based on physical location information from the TAIR database of the genome. Gene duplication in the *Arabidopsis* genome was analyzed with Multiple Collinearity Scan toolkit (MCScanX) (Wang et al., 2012), and if an *ADF* has more than one transcript, only the longest in the annotation was used. The collinearity relationship between *ADF* gene family members in *Arabidopsis* (dicot), *Glycine max* (dicot), *Zea mays* (monocot) and *Oryza sativa* (monocot) were shown using TBtools (Chen et al., 2020). The BLASTP (Altschul et al., 1990) was used with -evalue 1e-5 and MCScanX was used with MATCH_SIZE as 2. The calculation of Ka and Ks substitution of each duplicated *ADF* genes were performed using add_kaks_to_synteny.pl from MCScanX (Dataset S3).

### Physicochemical properties and protein location prediction

The physicochemical properties of the ADF, which are MW (kDa), theoretical pI, instability index, and GRAVY, were predicted by Expasy (https://web.expasy.org/protparam/). The subcellular localization of the ADF proteins were collected from TAIR, ThaleMine (https://bar.utoronto.ca/thalemine/begin.do), and predicted by WoLF PSORT(Horton et al., 2007).

### Phylogenetic tree construction of *ADF* isoforms

Sequences of ADF isoform proteins were used to build the phylogenetic tree, and the tree was constructed using the maximum likelihood (ML) method in RAxML (Randomized Axelerated Maximum Likelihood) (Stamatakis, 2014), with 1,00 bootstrap replicates. ML phylogenies tree was inferred and visualized by FigTree v1.4.4 (http://tree.bio.ed.ac.uk/software/figtree/).

### Single-cell RNA-sequencing (scRNA-seq) expression pattern of *Arabidopsis ADF* genes

A total of 34 single-cell dataset from 169 experiments from *Arabidopsis* in scPlantDB (He et al., 2024) was used to extracted the tissue and cell type information (Table 2). A total of 6 datasets, all Col-0 WT genotype under normal condition treatment and representing different tissues, were used to illustrate the *ADF* gene expression pattern, as shown in Fig. 6 and Dataset S6. For pollen tissue, only the Col-0 WT genotype from SRP374045 was selected to show the expression of *ADF7* and *ADF10*.

**Table 1.** *AtADF* gene family protein properties.

| Name | Chr | Position | Longest Isoform | Protein Characteristics |  |  |  |  | Subcellular Location |
| --- | --- | --- | --- | --- | --- | --- | --- | --- | --- |
|  |  |  |  | Amino Acids (aa) | MW (Kda) | Theoretical pI | Aliphatic index | GRAVY |  |
| AtADF1<br>(AT3G46010) | Chr 3 | 16909217-16910845 | AT3G4601<br>0.2 | 150 | 17.339 | 7.62 | 80.67 | -0.289 | Cytoplasm;<br>Chloroplast;<br>vascular tissues<br>of all organs |
| AtADF2<br>(AT3G46000) | Chr 3 | 16907334-16909119 | AT3G4600<br>0.2 | 156 | 17.942 | 6.59 | 74.42 | -0.362 | Cytoplasm; |
| AtADF3<br>(AT5G59880) | Chr 5 | 24120164-24121949 | AT5G5988<br>0.1 | 139 | 15.922 | 5.94 | 75.83 | -0.415 | Cytoplasm |
| AtADF4<br>(AT5G59890) | Chr 5 | 24122348-24123912 | AT5G5989<br>0.1 | 139 | 16.034 | 6.14 | 83.53 | -0.313 | Cytoplasm |
| AtADF5<br>(AT2G16700) | Chr 2 | 7244500-7246069 | AT2G1670<br>0.1 | 143 | 16.467 | 6.83 | 70.91 | -0.348 | Cytoplasm;<br>Expressed<br>exclusively in root tip<br>meristem. |
| AtADF6<br>(AT2G31200) | Chr<br>2 | 13293953-13295410 | AT2G3120<br>0.1 | 146 | 16.70<br>7 | 5.46 | 70.0<br>7 | -<br>0.57<br>4 | Cytoplasm;<br>Expressed in<br>vascular tissues<br>of all organs. |
| AtADF7<br>(AT4G25590) | Chr<br>4 | 13058936-13060250 | AT4G2559<br>0.1 | 137 | 15.82<br>0 | 5.08 | 73.3<br>6 | -<br>0.38<br>9 | Nucleus;<br>Specifically<br>expressed in<br>pollen. |
| AtADF8<br>(AT4G00680) | Chr<br>4 | 279596-281072 | AT4G0068<br>0.2 | 150 | 17.35<br>3 | 5.12 | 71.4<br>7 | -<br>0.46<br>1 | Cytoplasm;<br>Expressed in the<br>root trichoblast<br>cells and<br>developed root<br>hairs. |
| AtADF9<br>(AT4G34970) | Chr<br>4 | 16653704-16654955 | AT4G3497<br>0.1 | 141 | 16.36<br>9 | 7.66 | 60.8<br>5 | -<br>0.55<br>1 | Cytoplasm |
| AtADF10<br>(AT5G52360) | Chr<br>5 | 21257934-21259408 | AT5G5236<br>0.1 | 137 | 15.88<br>4 | 5.57 | 76.2<br>0 | -<br>0.44<br>5 | Nucleus;<br>Specifically<br>expressed in<br>pollen. |
| AtADF11<br>(AT1G01750) | Chr<br>1 | 275188-276310 | AT1G0175<br>0.2 | 145 | 16.88<br>6 | 5.54 | 72.6<br>2 | -<br>0.56<br>1 | Nucleus |
| ADF<br>(AT3G45990;<br>Putative) | Chr<br>3 | 16900778-16904136 | AT3G4599<br>0.1 | 133 | 15.90<br>4 | 7.74 | 79.9<br>2 | -<br>0.59<br>9 | Unknown |

**Table 2:**
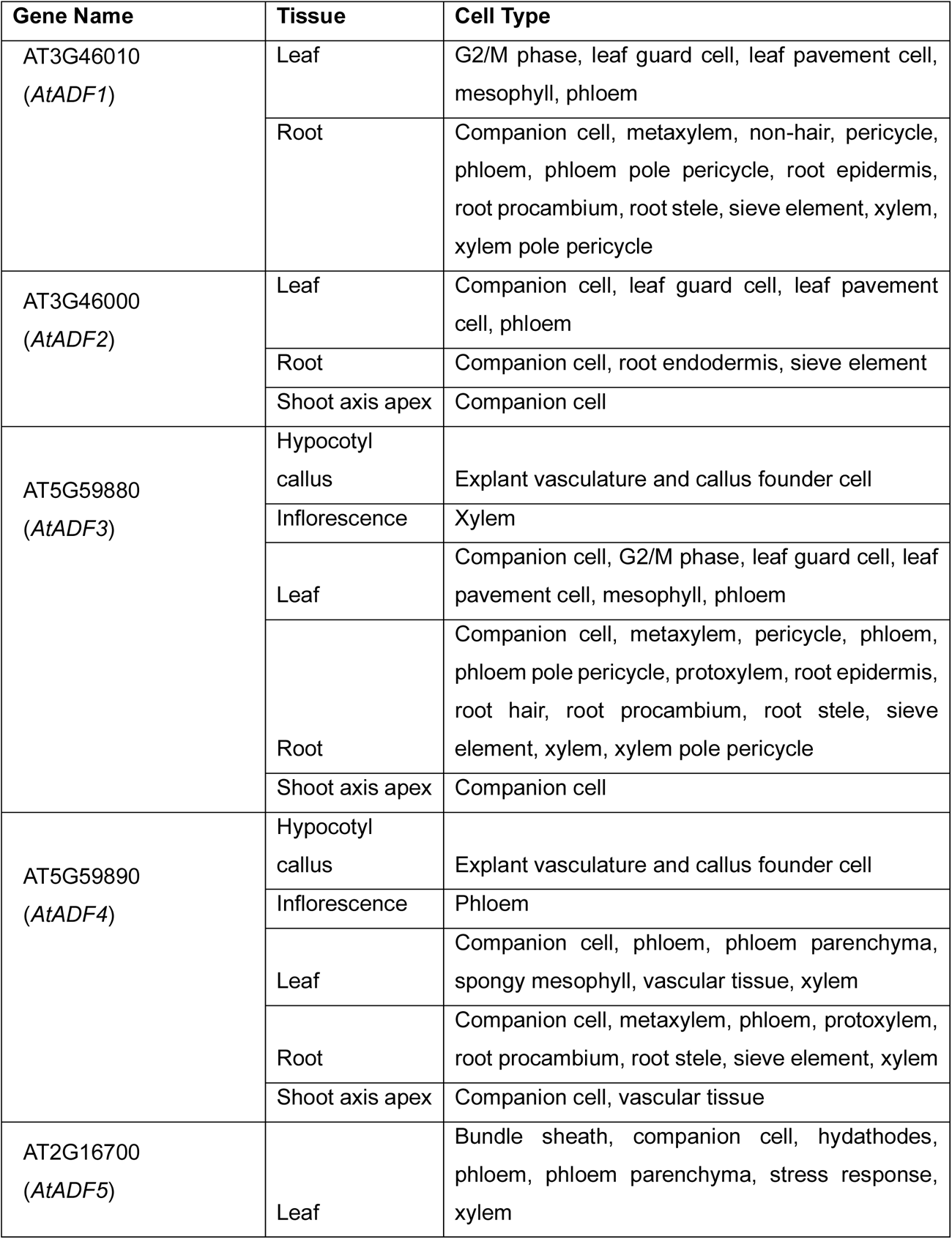
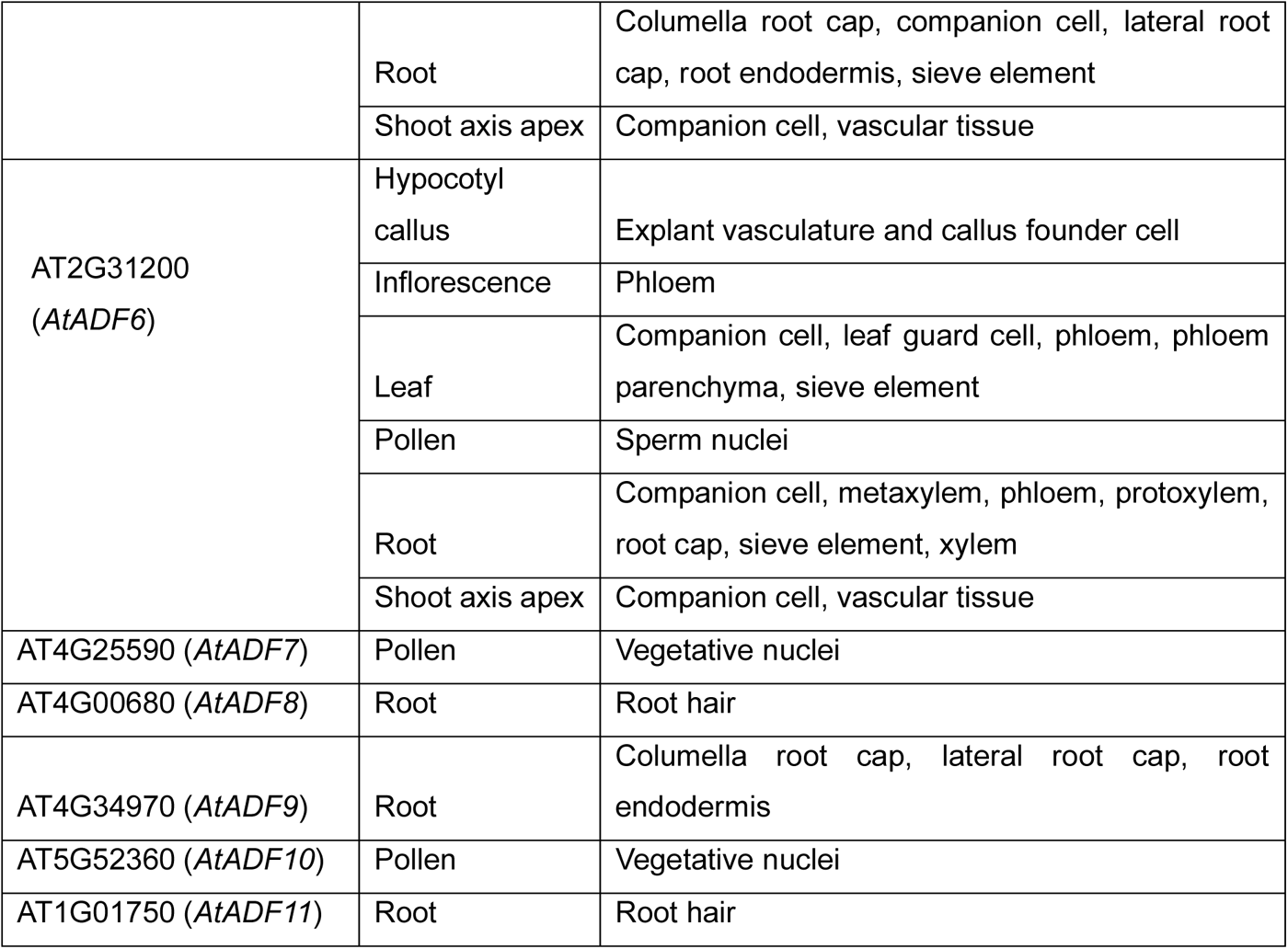
*AtADF* sing-cell gene expression in various tissue and cell type.

| Gene Name | Tissue | Cell Type |
| --- | --- | --- |
| AT3G46010<br>(*AtADF1*) | Leaf | G2/M phase, leaf guard cell, leaf pavement cell, mesophyll, phloem |
|  | Root | Companion cell, metaxylem, non-hair, pericycle, phloem, phloem pole pericycle, root epidermis, root procambium, root stele, sieve element, xylem, xylem pole pericycle |
| AT3G46000<br>(*AtADF2*) | Leaf | Companion cell, leaf guard cell, leaf pavement cell, phloem |
|  | Root | Companion cell, root endodermis, sieve element |
|  | Shoot axis apex | Companion cell |
| AT5G59880<br>(*AtADF3*) | Hypocotyl callus | Explant vasculature and callus founder cell |
|  | Inflorescence | Xylem |
|  | Leaf | Companion cell, G2/M phase, leaf guard cell, leaf pavement cell, mesophyll, phloem |
|  | Root | Companion cell, metaxylem, pericycle, phloem, phloem pole pericycle, protoxylem, root epidermis, root hair, root procambium, root stele, sieve element, xylem, xylem pole pericycle |
|  | Shoot axis apex | Companion cell |
| AT5G59890<br>(*AtADF4*) | Hypocotyl callus | Explant vasculature and callus founder cell |
|  | Inflorescence | Phloem |
|  | Leaf | Companion cell, phloem, phloem parenchyma, spongy mesophyll, vascular tissue, xylem |
|  | Root | Companion cell, metaxylem, phloem, protoxylem, root procambium, root stele, sieve element, xylem |
|  | Shoot axis apex | Companion cell, vascular tissue |
| AT2G16700<br>(*AtADF5*) | Leaf | Bundle sheath, companion cell, hydathodes, phloem, phloem parenchyma, stress response, xylem |
|  | Root | Columella root cap, companion cell, lateral root cap, root endodermis, sieve element |
|  | Shoot axis apex | Companion cell, vascular tissue |
| AT2G31200<br>(*AtADF6*) | Hypocotyl callus | Explant vasculature and callus founder cell |
|  | Inflorescence | Phloem |
|  | Leaf | Companion cell, leaf guard cell, phloem, phloem parenchyma, sieve element |
|  | Pollen | Sperm nuclei |
|  | Root | Companion cell, metaxylem, phloem, protoxylem, root cap, sieve element, xylem |
|  | Shoot axis apex | Companion cell, vascular tissue |
| AT4G25590 (*AtADF7*) | Pollen | Vegetative nuclei |
| AT4G00680 (*AtADF8*) | Root | Root hair |
| AT4G34970 (*AtADF9*) | Root | Columella root cap, lateral root cap, root endodermis |
| AT5G52360 (*AtADF10*) | Pollen | Vegetative nuclei |
| AT1G01750 (*AtADF11*) | Root | Root hair |

### Expression pattern of *ADF* genes in various tissues under different stresses

A total of 4,612 RNA-Seq sample with Col-0 WT genotype were selected to show the *ADF* isoform expression pattern in Fig. 7. These samples were chosen from the total of 24,123 RNA-Seq sample of *Arabidopsis*. TPM values of all *ADF* genes isoforms were used for doing clustering. The within-cluster sum of squares was calculated for K values ranged from 1 to 1000 using k-means, and the resulting plot of the within-cluster sum of squares against K was generated using R. The optimal K value was determined where the elbow value equals 0.8. The R package pheatmap was used to generate heatmaps for each cluster. The number of specific samples in each cluster is listed in Dataset S7.

## Results

### Prediction of potential immune-associated function genes

To predict genes from *Arabidopsis* with potential immune signaling function(s), the gene expression values of 24,123 RNA-Seq datasets and 34 single-cell datasets were evaluated based on the known identities of 3,107 immune genes possessing features known to associate with immunity/defense roles in plants. With this, we used this information to build gene models using four machine learning algorithms: k-nearest neighbors (KNN) (Dasarathy, 1991), decision tree (Breiman, 1984), random forest (Ho, 1995), and neural networks (NNs) (Hardesty, 2017) (Fig. S1). The accuracy for KKN, decision tree, random forest, and NNs were 0.69, 0.79, 0.89, and 0.83, respectively (Dataset S1). From this analysis, a total of 38 genes were predicted as having potential immune-related functions (Dataset S2).

Using these 38 gene candidates, we next explored their current known functions from TAIR, and the results showed some of them are involved in different biological process, such as response to salt, salicylic acid (SA), hypoxia, and photosynthesis, while the remaining candidates had unknown biological processes. As shown in Dataset S2, the promoter of gene candidate ATCG00280 contains a blue-light responsive element and is involved in the biological process of photosynthesis and light reaction. Recently research by Wu and Yang revealed that CRY1, a blue light photoreceptor, is involved in promoting R protein-mediated plant resistance through plant innate immunity (Wu and Yang, 2010). This infers that ATCG00280, a predicted potential immune-associated function genes candidate, is involved in plant innate immunity. Furthermore, this implies that other gene candidates involved in photosynthesis and photorespiration may also play a role in plant innate immunity. Additionally, gene candidate AT4G05320 is involved in SA signaling, which is a defense signal that activates plant immunity. This is important because the above results demonstrated that our model is robust in predicting potential immune-related functions genes and helpful for predicting the potential function of genes with very limited information, which could be helpful in narrowing down the targeting of new immune candidate genes for exploration in plants. In additionally, the identification of the immune genes enables researchers to better understand the immune response mechanisms and build a comprehensive picture of the highly dynamic in terms of temporal and spatial scales plant immune system.

Within the vast area of immune genes, one area of particular interest lies in the study of gene family that play crucial roles in actin cytoskeleton, promoting immune response among eukaryotes. Among these family, the *ADF* gene family stands out for its crucial role as actin-binding proteins to maintain the balance between F-actin and G-actin by producing filament ends conducive to new filament initiation. The ADF proteins is a conserved ancient protein family in all eukaryotes, and studies have shown that *ADFs* are involved in various biological process, including innate immune signaling. The gene expression pattern with *ADF* gene family are tissue-specific, with physiological and functional role differences. The *ADF* gene family shaped by the selective pressures during evolution, and understanding the evolution of *ADF* gene family is essential for comprehending how organisms defend themselves against dangers.

### Evolutionary relationships among *ADF* members in representative eukaryotic species

To investigate the evolutionary relationship(s) and characteristics of the *ADF* genes with other homologous genes, 38 eukaryotes species and 1 bacteria species – an outgroup – were used to construct a species tree (Fig. 1a) and using the longest amino acid sequences of ADF proteins from all species were used to construct an *ADF* gene family phylogenetic tree (Fig. 1b). As shown, phylogenetic analysis revealed that ADF most likely originated in the last common ancestor of animals, fungi, and plants. Two main ADF clades were classified as plant-specific and not plant-specific. The clade of not plant-specific integrates yeast *ADF* genes together with invertebrates, the protists, the metazoan, the fungi, and the vertebrates, which indicates that this is most likely the ancestral clade of the *ADF* gene family. Plant-specific clade only integrates plant-specific. Taken together, these findings suggest that plant-specific clade emerged in the last common ancestor of eukaryotes. The ADF in plant and other kingdoms are completely separated.

**Fig. 1.**
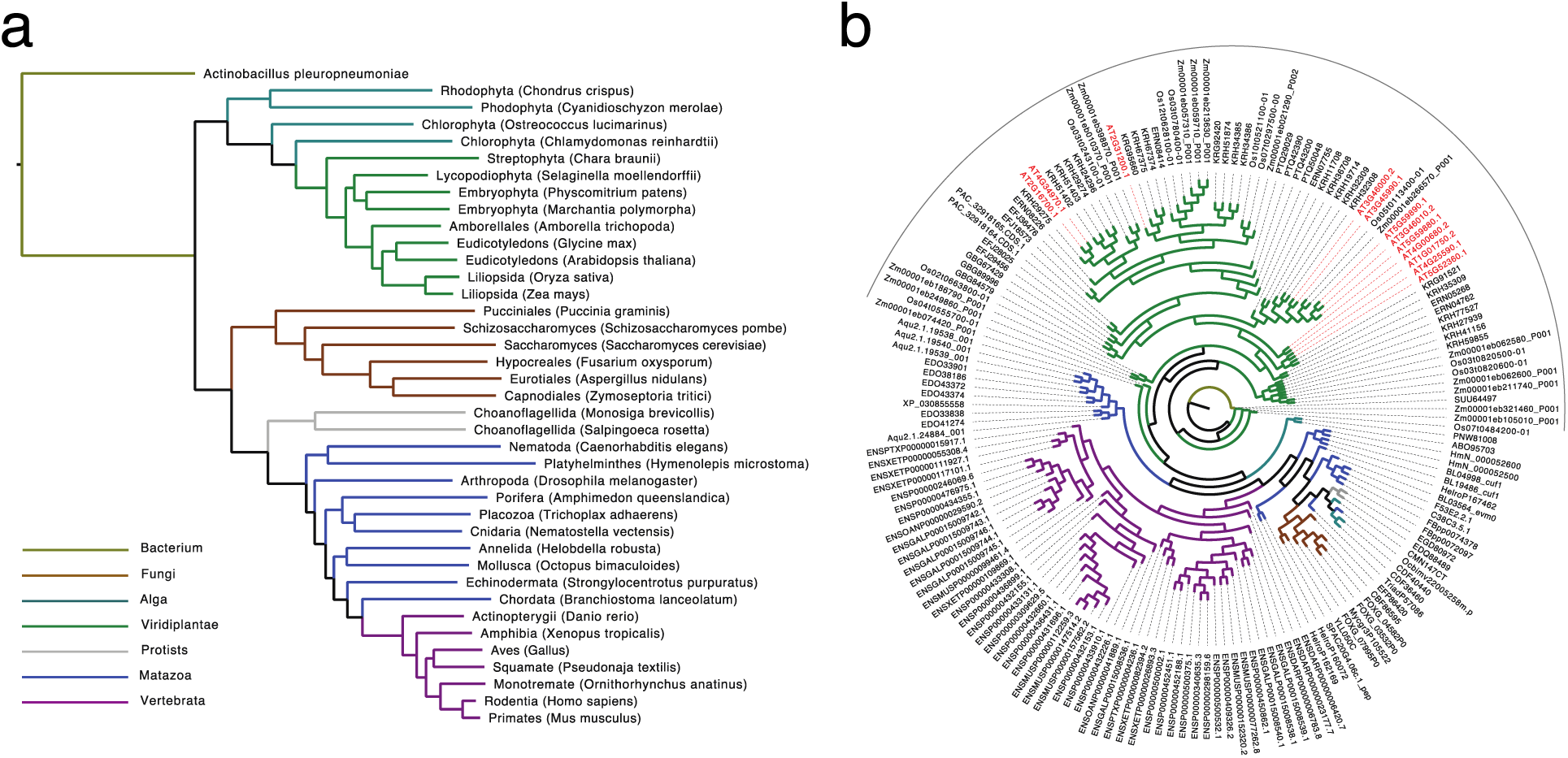
Species evolution tree and Phylogenetic tree of ADFs. **(a) Evolution relationship among 38 eukaryotes with 1 bacterium as outgroup.** (b) **Phylogenetic tree of ADF family members in the 38 eukaryotes species and most similar ADF protein in 1 bacterium by using the longest isoform.** ADF proteins in *Actinobacillus pleuropneumoniae, Amborella trichopoda, Amphimedon queenslandica, Arabidopsis thaliana, Aspergillus nidulans, Branchiostoma lanceolatum, Caenorhabditis elegans, Chara braunii, Chlamydomonas reinhardtii, Chondrus crispus, Cyanidioschyzon merolae, Danio rerio, Drosophila melanogaster, Fusarium oxysporum, Gallus, Glycine max, Helobdella robusta, Homo sapiens, Hymenolepis microstoma, Marchantia polymorpha, Monosiga brevicollis, Mus musculus, Nematostella vectensis, Octopus bimaculoides, Ornithorhynchus anatinus, Oryza sativa, Ostreococcus lucimarinus, Physcomitrium patens, Pseudonaja textilis, Puccinia graminis, Saccharomyces cerevisiae, Salpingoeca rosetta, Schizosaccharomyces pombe, Selaginella moellendorffii, Strongylocentrotus purpuratus, Trichoplax adhaerens, Xenopus tropicalis, Zea mays* and *Zymoseptoria tritici* are prefixed by SU, ER, Aq, AT, CB, BL, C3 or F5, GB, PN, CD, CM, ENSD, FB, FO, ENSG, KR, He, ENSP0, Hm, PT, EDQ, ENSM, EDO, Oc, ENSO, Os, AB, PA, ENSPT, EFP, YL, EG, SP, EFJ, XP, Tr, ENSX, Zm and My. ADF in *Arabidopsis thaliana* are marked in red. The grey half circle line indicates the plant-specific clade. ADF proteins of *Arabidopsis thaliana* are marked as red. The different color lines represent different kingdoms.

### Conserved motifs analysis of ADF family member in eight representative species

A species tree of eight representative species was constructed by OrthoFinder (Fig. 2). Based on this, we first identified the conserved motif(s) present within the *ADF* gene family. As shown in Fig. 2, we observed that all plant ADFs possess a unique motif (i.e., motif 7), with an additional motif (i.e., motif 4) present in all *Arabidopsis* funcational *ADF* genes. We hypothesis that this feature underpins the observation as to why all the ADFs in plants comprise a plant-specific clade, including why the single putative *ADF* from *Arabidopsis* is without function (Ruzicka et al., 2007). To investigate the ADF family members in plant, *ADFs* from *Arabidopsis* will be used to study evolution history and function prediction.

**Fig. 2.**
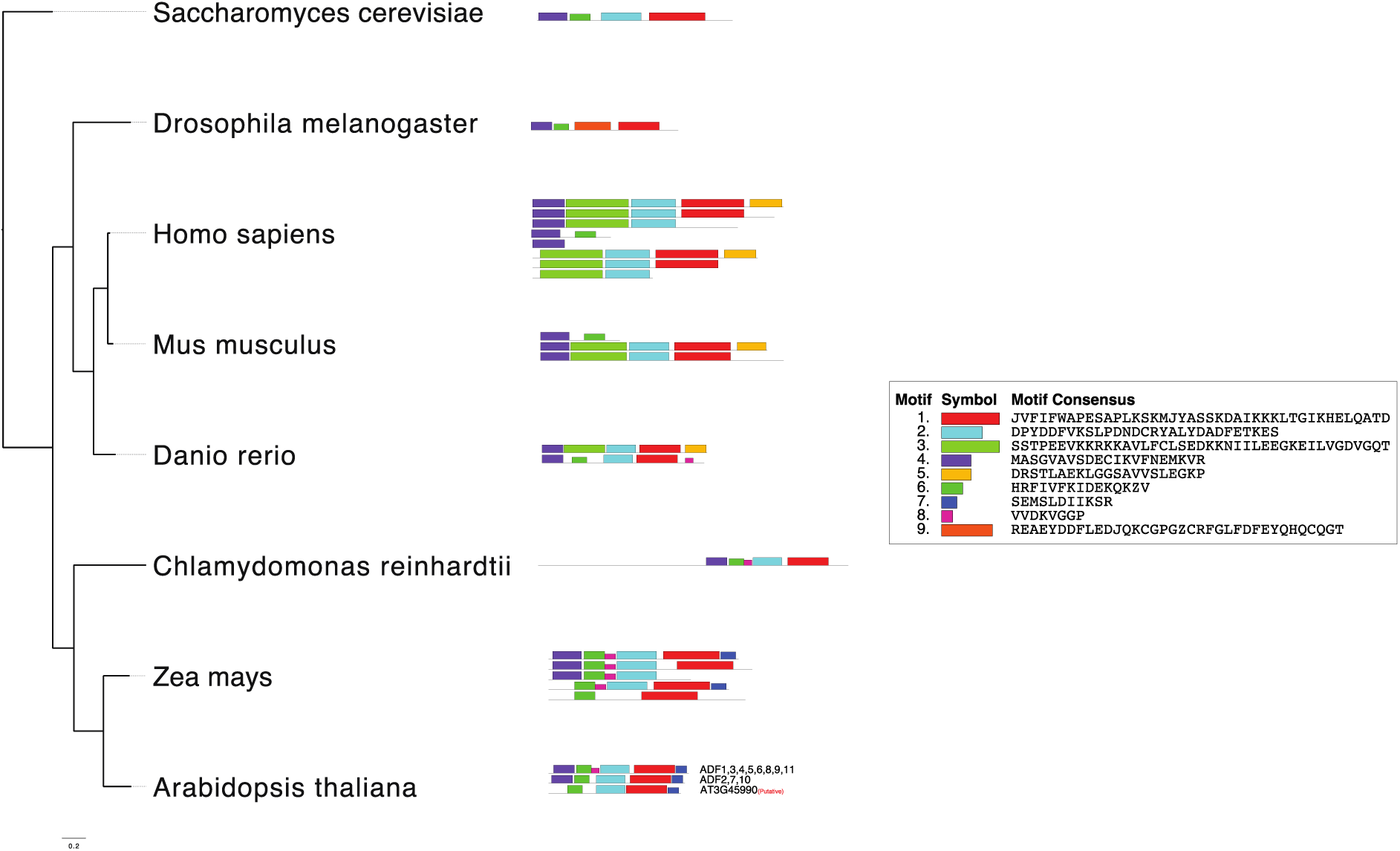
Phylogenetic relationship and motif composition of ADFs among eight representative species. (a) Phylogenetic relationship of ADFs. (b) Distribution of predicted conserved motifs type of ADFs.

### Cis-acting elements of the *ADF* gene family

Phylogenetic analysis of all ADF classifications indicates these 12 genes in *Arabidopsis* could be divided into three clusters, as shown in Fig. 3a, and all the *ADF* genes’ structure are relatively simple with only three exons as previously described (Inada, 2018). To extend this analysis, we next investigated the regulatory elements controlling ADF expression. Cis-acting elements are important for gene transcription. Here, a 1000 bp sequence upstream of the transcription start site of transcripts was extracted as promoters to predict cis-acting elements in 11 functional *Arabidopsis ADF* genes (Fig. 3b). A total of 22 cis-acting element types were identified and classified into five categories: cell cycle (2), development (1), hormone (1), stress (16), and transcription (3), for a total of 693 predicted cis-acting elements in *Arabidopsis*. Additionally, all *ADF* genes contain TATA-box and MYB, with the number of the TATA-box elements (293) being the greatest. The number of MYB and TATA-box was the highest among stress relative elements kinds, and the number of the development elements was the lowest (Dataset S3).

**Fig. 3.**
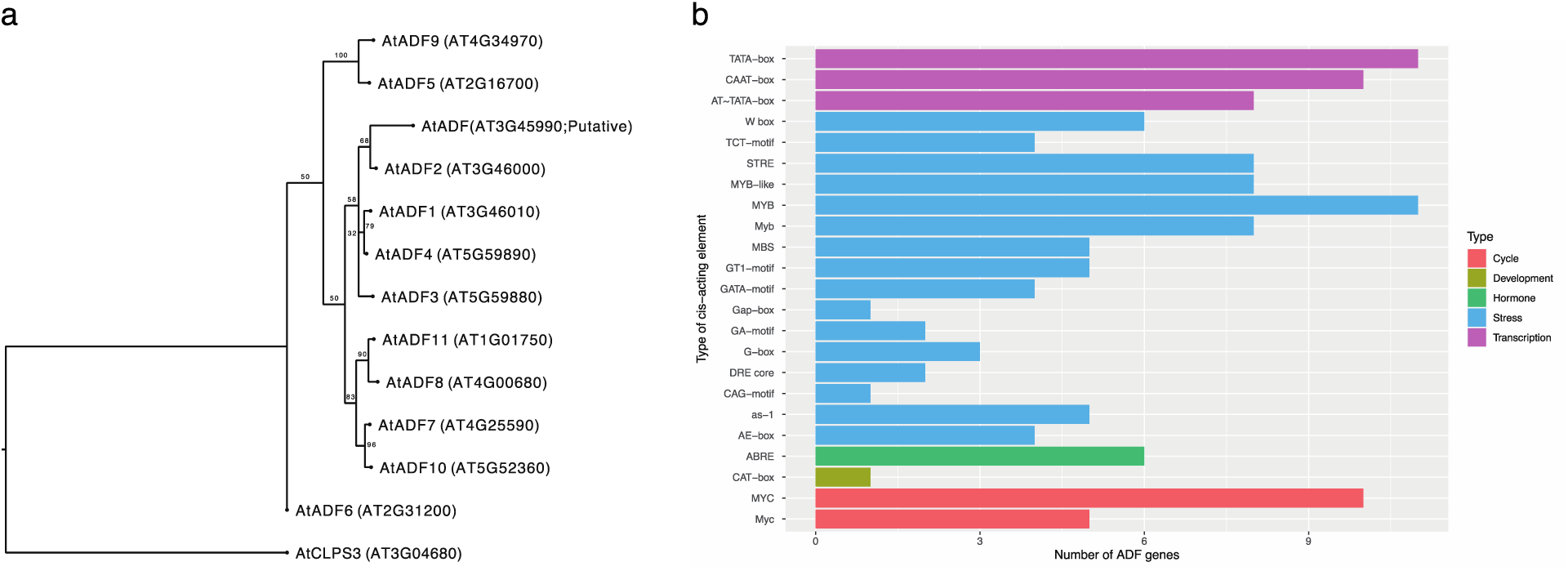
Phylogenetic relationship and prediction and analysis of cis-acting elements in the *AtADF* gene family promoter region. (a) The Maximum Likelihood phylogenetic tree of ADFs was constructed using the longest amino acid sequence from the longest isoform with CLPS3 as outgroup. (b) Classification cis-acting elements. Twenty-two types of cis-acting elements were divided into five categories. Different colors represent different categories.

Analysis of the physicochemical properties of proteins within the *Arabidopsis ADF* gene family Basic information about the *ADF* gene family and the proteins is shown in the Table 1. The longest genome sequence is *ADF* (AT3G45990; Putative), at 3,359bp, and the shortest is 945 bp (*ADF11*). The average length is 1,490 bp for all *ADFs*. Next, we analyzed the physicochemical properties of ADF proteins, including amino acid length, molecular weight (MW), theoretical pI, aliphatic index, and the grand average of hydropathy (GRAVY) (Table 1). The amino acids length of each ADF was found to be similar, ranging from 133 to 150. The molecular weight varied from 15,820 kDa (AtADF7) to 17.942 kDa (ADF2), while the GRAVY of all *ADF* genes were below zero. The maximum aliphatic index value was 83.53 (AtADF4), and the minimum value was 70.07 (AtADF6). The results of hydrophilicity and hydrophobicity analysis indicated that all ADF family proteins are hydrophilic proteins. The results of subcellular localization of the proteins analyzed using online software and information got from TAIR showed that all ADF family proteins are cytoplasmic proteins.

### Chromosome localization and collinearity analysis *ADF* genes family

The relationship between the location of *Arabidopsis ADF* gene family members on each chromosome and the collinearity within the *Arabidopsis* genome is shown in Fig. 4. A total of 12 *Arabidopsis ADF* genes – 11 functional 1 putative – are partially distributed on the chromosomes of *Arabidopsis* and are found on all five chromosomes. Specifically, only one *ADF* was located on chromosome 1 (*ADF11*), two on chromosome 2 (*ADF5* and *ADF6*), three genes on chromosome 3 (*ADF1*, *ADF2*, and *ADF* (putative)), three on chromosome 4 (*ADF7*, *ADF8,* and *ADF9*) and three on chromosome 5 (*ADF3*, *ADF4,* and *ADF10*). All *ADFs* are relatively close to the telomeric region of the chromosome. The above results suggest that ADF gens have undergone evolutionary conservation or specific adaptation, and genes within *ADF* gene family may have different gene expression and function due to the instability of the telomeric region of the chromosome. This is consistent with previous research showing that *ADF* members have distinct tissue-specific patterns in both physiological and functional terms (Wang et al., 2023, Miklis et al., 2007, Porter et al., 2012, Peng and Huang, 2006).

**Fig. 4.**
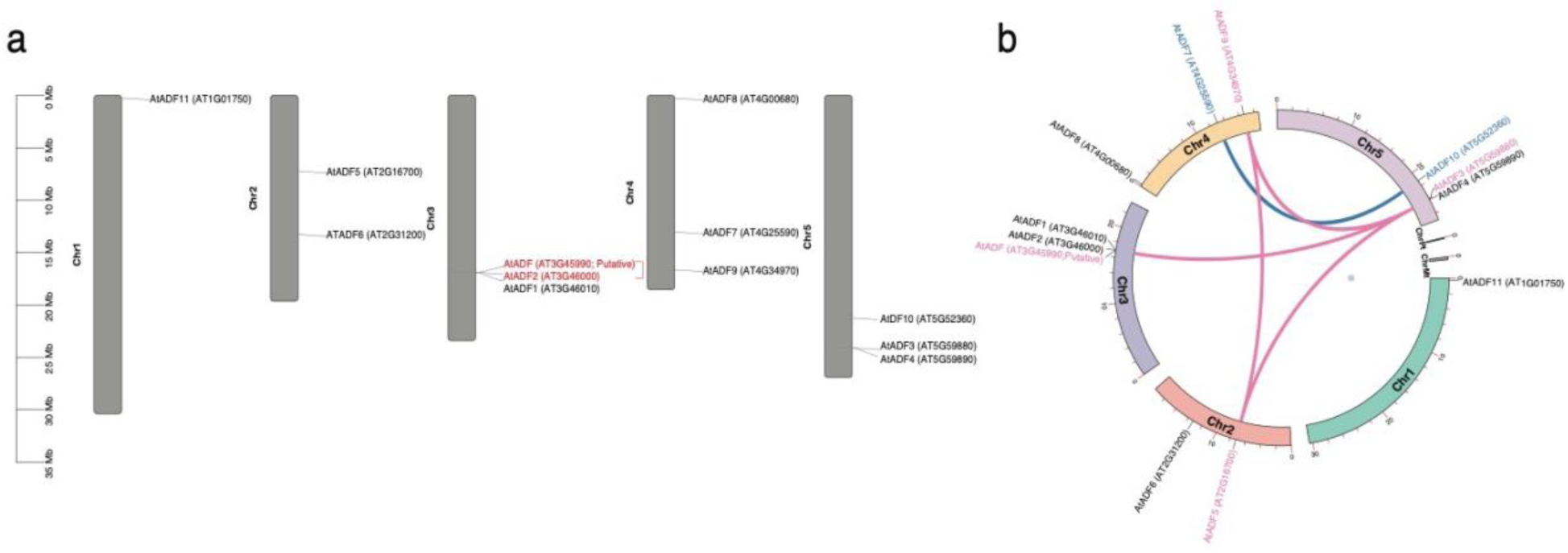
Chromosomal location and duplication events of ADF in *Arabidopsis*. (a) Chromosomal location of *ADF* genes in *Arabidopsis*. Twelve *ADF* genes were distributed on chromosomes 1, 2, 3, 4 and 5. The chromosome numbers are indicated at the left side of each vertical bar. Red line between genes indicates tandem duplication of *ADF* genes. (b) Gene duplication events of AtADFs. The different colors represent seven chromosomes. The segmentally duplicated genes are linked by different color lines.

To determine the replication relationship(s) between genes within *Arabidopsis*, we next conducted a collinearity analysis. As shown in Fig 2b, we identified the presence of one tandem duplication between *ADF2* and the putative *ADF* gene. *ADF* gene family members have five segmental duplications relationships within species, which are *ADF3:ADF5*, *ADF3:AtADF9*, *ADF3:ADF* putative gene, *ADF5*:*ADF9* and *ADF7:ADF10*. The selection pressure on the *ADF* gene duplication events in *Arabidopsis* were evaluated by calculating the rates of non-synonymous (Ka) and synonymous (Ks) nucleotide substitutions (Dataset S4). The average Ka/Ks ratios for the five duplicated pairs were less than 1.00, implying the *ADF* genes evolved under strong purifying selection.

### Systematic evolutionary relationships among *ADF* gene members in monocots and dicots

To determine the phylogenetic mechanism of these 12 *ADF* genes of *Arabidopsis*, we examined the synteny between *Arabidopsis* and other three species, including *Glycine max* (dicot), *Zea mays* (monocot), and *Oryza sativa* (monocot). As shown in Fig. 5, *Arabidopsis* (dicot) had 12 collinearity with *Glycine max*, 1 collinearity with *Zea mays*, and 4 collinearities with *Oryza sativa*, respectively. *ADF1*, *ADF2*, *ADF4* and *ADF8* in *Arabidopsis* are without collinearity with genes from other three species (Dataset S5), while 1 collinearity pair was identified in all these four species. However, other orthologous gene pairs were only identified between two of them.

**Fig. 5.**
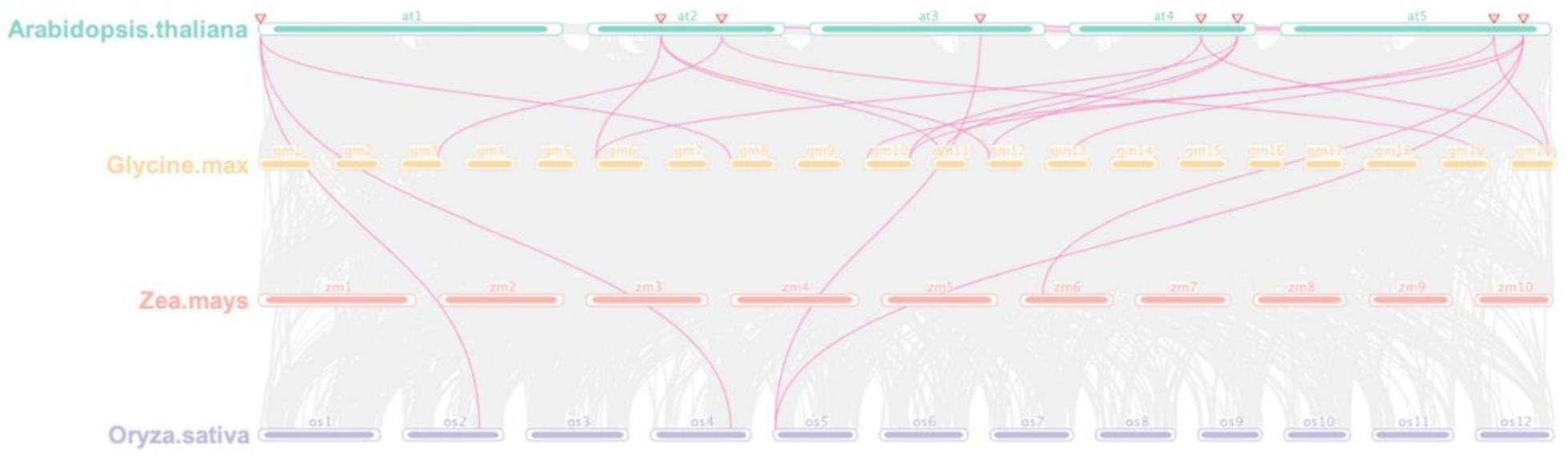
Collinearity relationship between *ADF* gene family members in *Arabidopsis thaliana* (dicto), *Glycine max* (dicot), *Zea mays* (monocot) and *Oryza sativa* (monocot). The chromosome number is marked above the chromosome. The chromosomes of *Arabidopsis thaliana*, *Glycine max*, *Zea mays* and *Oryza sativa* are marked with different colors. The collinear relationship between the *ADF* gene family members among four different species is connected by purple-colored lines. The red triangle represents the location of *ADF* genes on *Arabidopsis thaliana*.

**Fig. 6.**
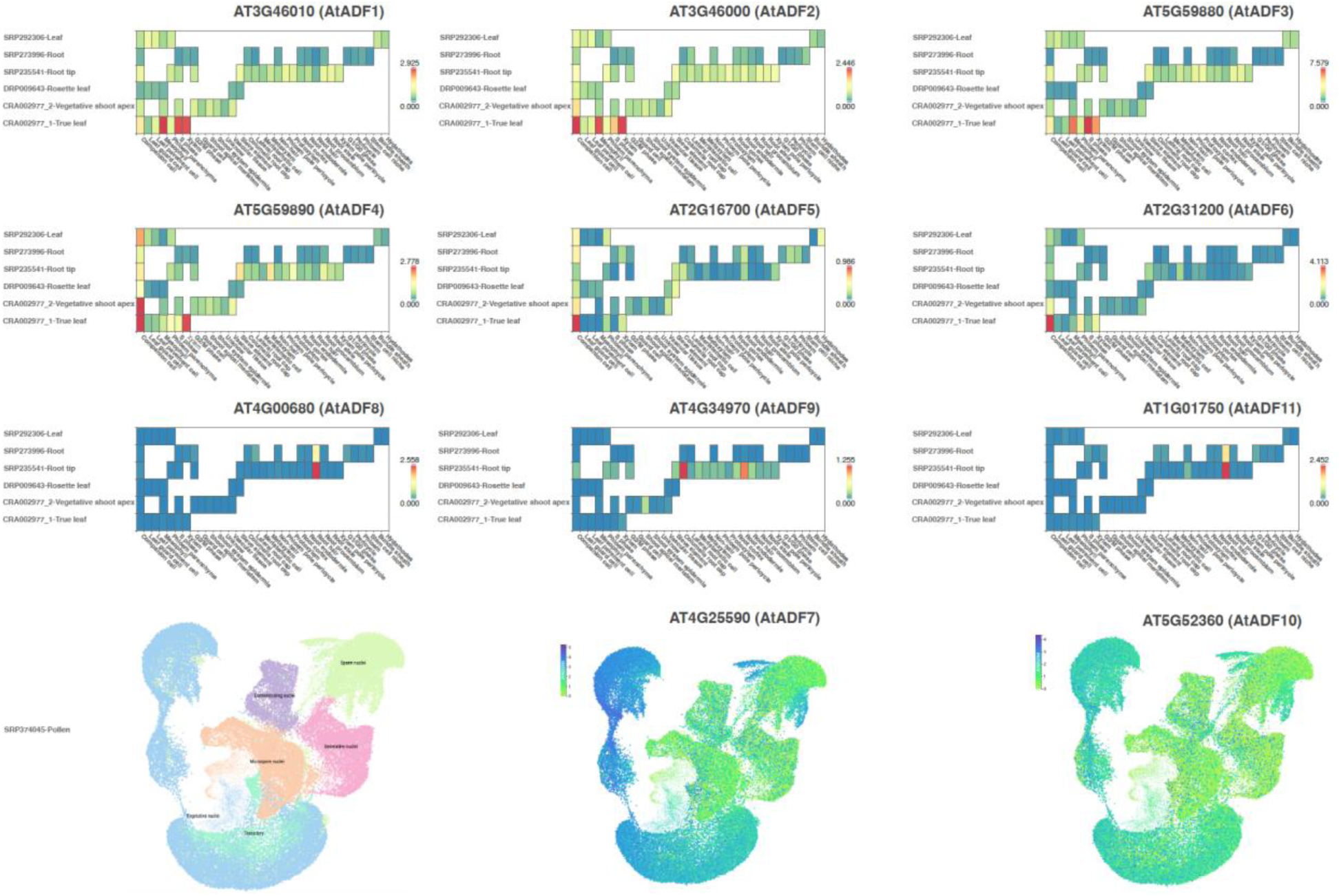
Single-Cell expression of *AtADF* genes in various tissue. Heatmap of transcript expression of AtADF. X axis is the cell-type cluster. Y axis is the tissue type. Red and blue indicate high and low expression values, respectively.

**Fig. 7.**
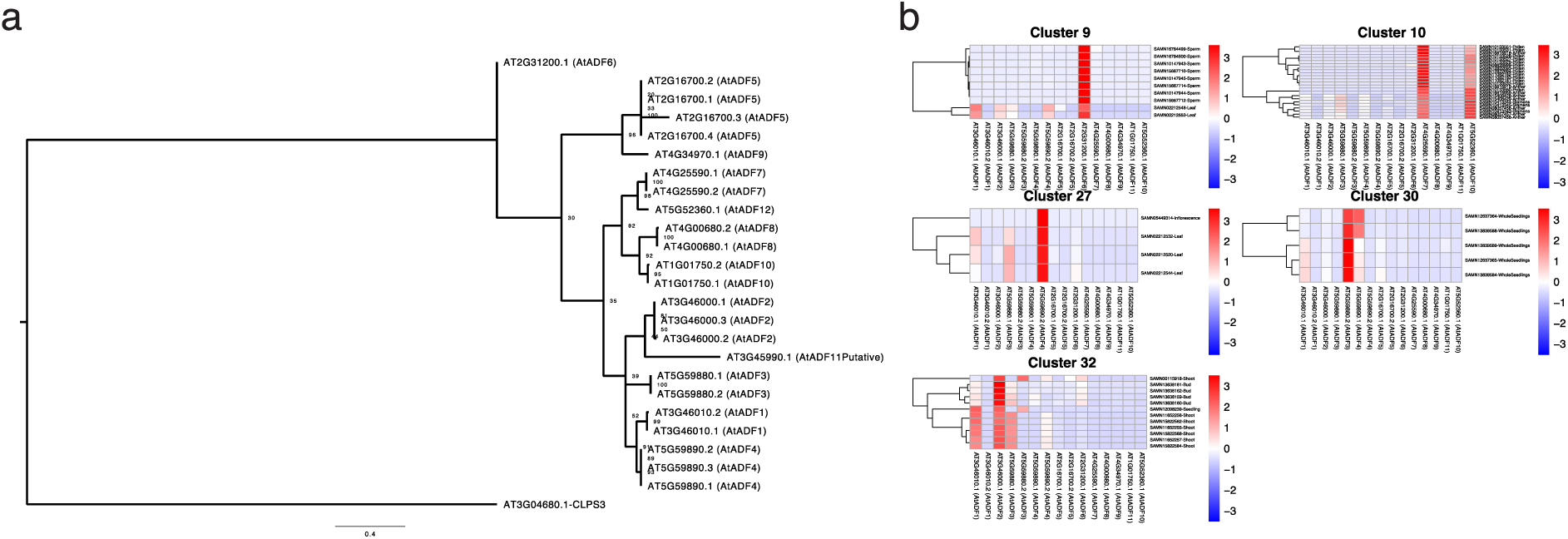
Phylogenetic relationship of *AtADF* gene isoform and RNA-Seq expression of *AtADF* genes in five not common clusters. (a) Phylogenetic relationship of *AtADFs* gene isoform. (b) Heatmap of expression of TPM value of AtADF isoform. Red and blue indicate high and low expression values, respectively.

The above results indicates that *ADF* gene family members in *Arabidopsis* are most closely related to those in *Glycine max*, likely due to their shared dicotyledonous ancestry and evolutionary history, which differs from the monocotyledonous species *Zea mays* and *Oryza sativa*. The 1 collinearity pair in all these four species and 4 *ADF* in *Arabidopsis* without collinearity with genes from other three species suggest that *ADF* orthologous pair was relatively conserved during evolution, but particular *ADFs* in *Arabidopsis* have undergone significant divergence or rearrangements, which may due to the genome assembly or gene deletion or chromosomal recombination occurred during evolution. The syntenic relationship analysis of *Arabidopsis ADF* gene family with other plant species can help provide the foundation of genetic relationship and gene function of species.

### Single-cell expression of *Arabidopsis ADF* in different tissues and cell types

The result of gene expression in single-cell RNA-Seq (scRNA-Seq) are shown in Fig. 6 and Table 2. The gene expression tissue and cell type information were extracted from scPlantDB (He et al., 2024). All of the 11 functional *ADF* gene were expressed in *Arabidopsis*, both *ADF7* and *ADF10* were only expressed in vegetative nuclei of pollen, which is consistent with results from previous research that they are specifically expressed in pollen (Bou Daher et al., 2011, Daher and Geitmann, 2012). *ADF1-6* were expressed in phloem cell of leaf, companion cell and sieve element cell in root. *ADF2-5* were expressed in companion cell of shoot axis apex. Additionally, *ADF3*, *4* and *6* were expressed in explant vasculature and callus founder cell of hypocotyl callus. Furthermore, *ADF8* and *ADF11* were expressed in root hair cell in root, while *ADF9* was expressed in columella root cap, lateral root cap and root endodermis. The above results are consistent with results from RNA-Seq, which indicates that *ADF1*,*ADF4,* and *ADF6* may play multiple roles in different tissue types compared to other *ADF* genes within of *Arabidopsis*.

### Gene expression pattern evolution of *Arabidopsis ADF* gene family

The gene expression clustering analysis grouped the genes into 33 distinct clusters as shown in Fig. S2. A total of 28 clusters (Cluster 1-8, 11-26, 28-29, 31 and 33) exhibit the same *ADF* isoform expression pattern, where *ADF3* (AT5G59880.1) is the most strongly expressed, and the other *ADF* isoforms show similar expression patterns across different tissues and treatments.

However, as shown in Fig. 7, the isoform expression pattern of *ADF* is unique in 5 clusters (Cluster 9, 10, 27, 30 and 32). Cluster 9 exhibited strong expression of *ADF6* (AT2G31200.1) in sperm and leaf, consistent with the scRNA-Seq results that *ADF6* are strongly expressed in companion cells within the leaf (Fig. 6). Cluster 10 showed strongly expression of *ADF7* (AT4G25590.1) and *ADF10* (AT5G52360.1) in pollen and anther, while cluster 27 exhibited has relative high expression of *ADF4* (AT5G59890.2) in leaf and inflorescence, which is also constant with the scRNA-Seq results that *ADF4* is strongly expressed in companion cell and xylem cell in leaf tissue (Fig. 6). Cluster 30 exhibited strong expression of *ADF3* (AT5G59880.2) in whole seedling, while the scRNA-seq also showed *ADF3* has relatively higher expression in various tissues, including leaf, root, root tip, rosette leaf, vegetative shoot apex and true leaf. Cluster 32 showed that *ADF2* (AT3G46000.1) has strong expression in shoot, bud, and seedlings. These data indicate that *ADF2*, *ADF3*, *ADF4*, *ADF6*, *ADF7,* and *ADF10* evolved new sub-functions over evolution, such as response to Fe, copper-deficiency response, and ABA signaling.

## Discussion

The completion of the *Arabidopsis* genome sequence provided the foundation for the identification of protein-coding gene (Arabidopsis Genome, 2000), and numerous individual gene studies, which have benefited from sequencing technologies and various available computational biology approaches, have provided foundational support for predicting gene category (Aromolaran et al., 2022). In the current study, to expand our knowledge of potential immune genes, we focused on developing and applying computational big data-driven approaches to predicate the potential immune genes. In total, our model predicted 38 potential immune-associated function genes in *Arabidopsis*. Among these gene cantates, we identified gene AT4G05320 is involved in SA signaling, which is a defense signal that activates plant immunity. We also identified gene ATCG00280, whose promoter contains a blue-light responsive element similar to that of the known known CRY1 gene (Wu and Yang, 2010) is involved in promoting R protein-mediated plant resistance through plant innate immunity as a blue light photoreceptor.

Previous research has shown that actin cytoskeleton plays a central role in various cellular functions and multiple cellular processes in eukaryotes, which is achieved through highly organized and highly dynamic polymerizing and depolymerizing with ABPs within plant cells, thereby improving plant immunity (Henty-Ridilla et al., 2013, Staiger and Blanchoin, 2006, Pollard and Cooper, 2009, Hussey et al., 2006, Szymanski and Cosgrove, 2009, Day et al., 2011, Staiger, 2000, Zhao et al., 2011). ADFs, as an ancient and conserved protein in all eukaryotes, are an important ABP that maintains the balance of F- and G-actin by depolymerizing and polymerizing, thereby increasing the dynamics of the actin in cytoskeleton and improve immunity (Andrianantoandro and Pollard, 2006, Pavlov et al., 2007).

In our study, we used 38 species to investigate the evolutionary relationships of *ADF* genes in eukaryotes. Our phylogenetic tree results showed that *ADF* most likely originated in the last common ancestor of animals, fungi, and plants, and that plant *ADFs* are completely separated from those in other kingdoms. The recent studies have also shown that *ADF* gene family is conserved and relatively small in higher plants (McCurdy et al., 2001, Feng et al., 2006, Roy-Zokan et al., 2015, Khatun et al., 2016, Huang et al., 2020). Additionally, we used eight representative species to explore the possible reasons why plants have a plant-specific clade within ADF gene family. The results showed that plant *ADFs* possess the unique motif (motif 7) maybe the reason, which may be the reason for the plant-specific clade. Interestingly, the research described herein supports the hypothesis that the plant *ADF* gene family is unique and likely emerged in the last common ancestor of eukaryotes.

Evolution is essential for species. Evolutionary processes lead to physiological or morphological differences between species, and gene duplication and loss contribute to the major difference of gene number between different species (Holland et al., 2017). Therefore, understanding the evolution history of a gene family is necessary for its study. In our study, the evolutionary relationships of *ADF* revealed that plants have a plant-specific clade that emerged in the last common ancestor of all eukaryotes. The evolutionary relationship of *ADF* between monocots and dicots revealed that one orthologous pair was relatively well conserved, while specific *ADFs* (*ADF1*, *2*,*4* and *8*) have undergone significant divergence or rearrangements. The reason for this may be due to the genome assembly, gene deletion, or chromosomal recombination occurred during evolution leading to differences in gene number between different species.

*Arabidopsis* is the model system of choice for research on plant and crop reproductive development due to its relatively simple genome (Meinke et al., 1998). Moreover, it is also an common and ideal species for studying gene family evolution and novel functions that have evolved, as most genes within its genome are members of gene families and massive available data from public databases, such as TAIR (https://www.arabidopsis.org/) and GEO (Cannon et al., 2004), can be utilized to support such studies. Segmental and tandem duplications are the major driving force behind gene family expansions during the evolution in *Arabidopsis* (Cannon et al., 2004). Consistent with this, our study detected one tandem duplication and five segmental duplications within *ADF* gene family members in *Arabidopsis*.

Indeed, if we track whole genome duplication events, genes in the same gene family originally came from a common ancestor through duplication, which is why all *ADF* genes has three exons and a relatively simple gene structure within *Arabidopsis* (Inada, 2018). This conserved gene structure is likely a results of strong purifying selection, which is supported by our analysis of Ka/Ks ratios showing that the *ADF* genes evolved under strong purifying selection during evolution. All these indicates that *ADF* genes family likely plays important roles in biological processes.

Gene expression patterns in various tissues can be used to reflect the different biological functions and evolutional patterns of genes within a gene family. In our study, we utilized the gene expression results from Col-0 WT *Arabidopsis* under various treatments, conducted clustering analysis and investigated the gene expression patterns in single-cell data. The results indicated that *ADF3* (AT5G59880.1) showed the highest expression among *ADF* genes across various tissues and under different conditions, and *ADF1,2* and *4* showed the second-highest relative expression, as shown in Fig. S2. Our result is consistent with recent research that has shown *ADF1-4* are expressed throughout plants at relatively high levels within *Arabidopsis*, indicating their importance in plant biology (Ruzicka et al., 2007). Recent research identified that *ADF7* and *ADF10* are only expressed in flowers, particularly in pollens (MacRobbie and Kurup, 2007), which is also consistent with our results showing these ADFs are strongly expressed in pollen and anther (Fig. 3a, Fig. 6 and Fig. 7).

The previous research has shown that some genes within a gene family retain the same function as their common ancestor, while others evolve novel sub-functions after gene expansion, and some may become pseudogenes (Nan et al., 2017). Our results showed *ADF7* and *ADF10,* which arose through segmental duplication, are in in the same clade and have retained the same function (Fig. 3a, Fig. 4b, Fig. 6 and Fig. 7), suggesting that duplication genes can retain the same function as their common ancestral genes. However, *ADF5* and *ADF9* are in the same clade (Fig. 3a), but *ADF9* has noticeably lower expression levels compared to *ADF5* in companion cells, phloem parenchyma and hydathodes in leaf, companion cells in root, indicating that duplication genes can also lose the function of their ancestral genes (Fig. 6). However, the expression level of *ADF9* in columella root cap is significantly higher than *ADF5* (Fig. 6), indicating that duplication genes can also evolve novel sub-functions beyond those of their ancestral genes. Compared to previous research, our expression pattern of *ADF* from transcript levels indicates that *ADF1*, *ADF4* and *ADF6* may play the multiple roles in different tissues compared to other *ADF* genes (Fig. 6), and *ADF2*, *ADF3, ADF4*, *ADF6*, *ADF7* and *ADF10* have evolved new sub-function for responding to stresses within *Arabidopsis* (Fig. 7).

*Arabidopsis* is one of the most important model plants for crop reproductive development. The *ADF* gene family has been reported to play a crucial role in plant development and stress response. Although several evolutionary function prediction studies have been conducted for the *ADF* gene family in *Arabidopsis*, no study has utilized big data-driven computational approaches study the evolution of this gene family and the evolution of their gene function.

However, with the increasing number of computational and data-rich resources available for *Arabidopsis*, combined with advances in computational biology, informatics-driven prediction of *ADF* gene function has become more feasible and powerful, guiding experimental studies like the work described herein. The potential immune-associated function genes predicted in our study could be used as important candidate genes for stress resistance research using the immune system for plant, and our findings contribute to the foundation knowledge for functional research on *ADF* genes, providing a valuable reference for stress-resistant breeding of crops.

While many immune gene families, such as ADF, evolved novel sub-function of each of member during evolution.

## Acknowledgments

We would like to acknowledge the help from Thilanka Ranaweera and Dr. Shin-Han Shiu for advice during the development of the research, and the help from Dr. Pai Li for providing meaningful background information of *ADF* gene family.

## Author contributions

HC and BD planned and designed the research. HC analyzed data. HC and BD wrote the papers. All authors discussed the results and approved the final manuscript for publication.

### Data availability

The codes for immune gene prediction and the gene expression of 24,123 RNA-Seq datasets, are pending at the following address https://github.com/chenh9313/Sup_ADF.git.

## Supplementary Figures

**Fig. S1.**
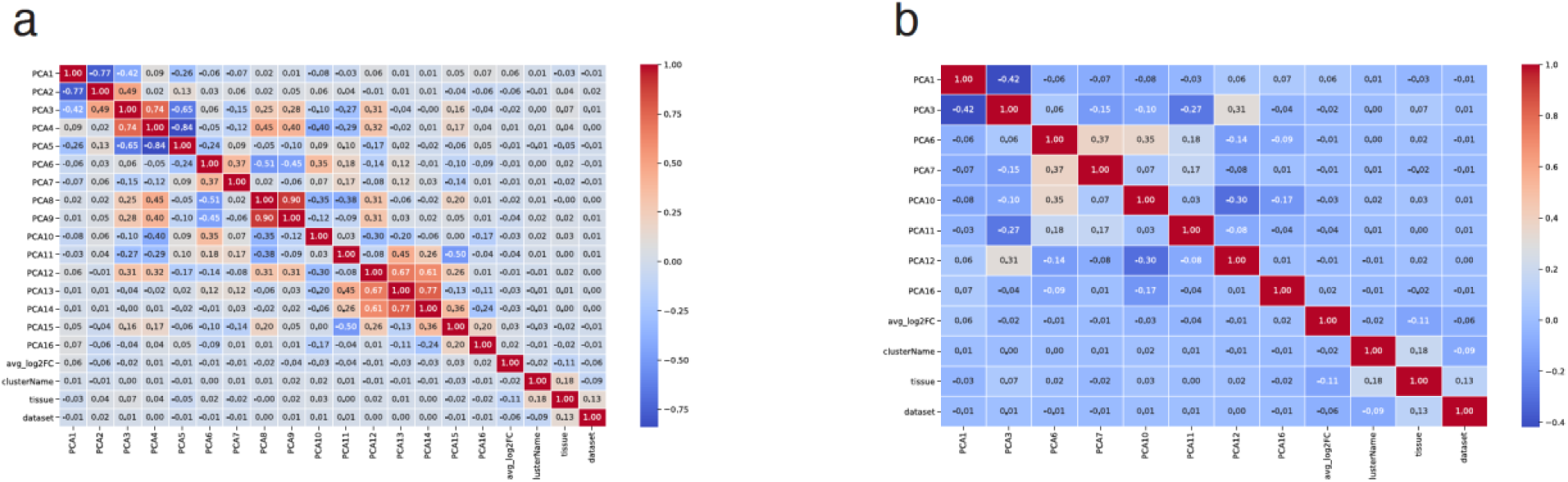
Correlation calculation between features. (a) Correlation calculation between all the features. (b) Correlation calculation between all the features with the correlation score <= 0.5.

**Fig. S2.**
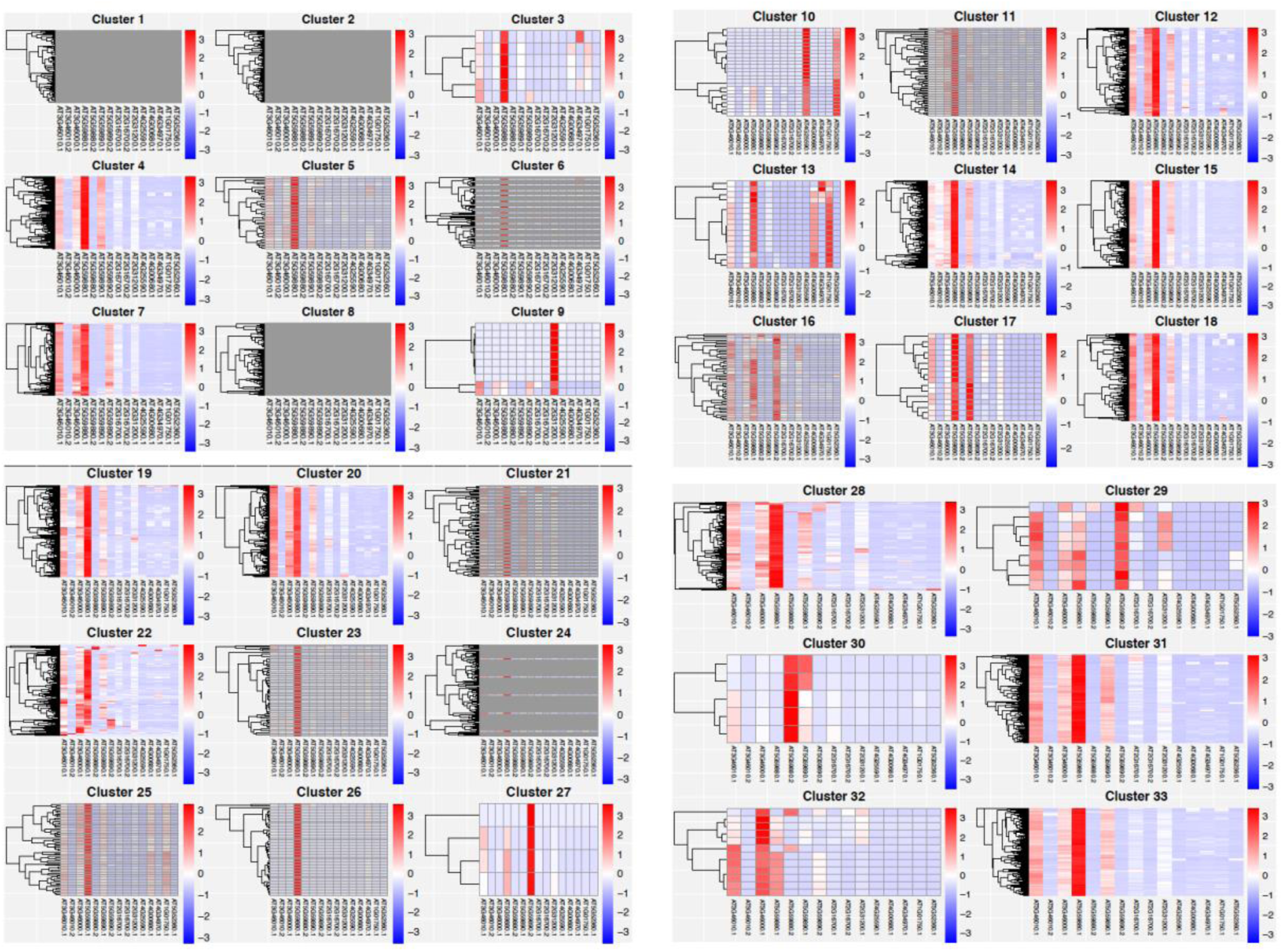
Total of 33 clusters of gene expression pattern.

## References

Altschul, S. F., Gish, W., Miller, W., Myers, E. W. & Lipman, D. J. 1990. Basic local alignment search tool. J Mol Biol, 215, 403–10.

Altschul, S. F., Madden, T. L., Schaffer, A. A., Zhang, J., Zhang, Z., Miller, W. & Lipman, D. J. 1997. Gapped BLAST and PSI-BLAST: a new generation of protein database search programs. Nucleic Acids Res, 25, 3389–402.

Andrianantoandro, E. & Pollard, T. D. 2006. Mechanism of actin filament turnover by severing and nucleation at different concentrations of ADF/cofilin. Mol Cell, 24, 13–23.

ARABIDOPSIS Genome, I. 2000. Analysis of the genome sequence of the flowering plant Arabidopsis thaliana. Nature, 408, 796–815.

Aromolaran, O., Aromolaran, D., Isewon, I. & Oyelade, J. 2022. Corrigendum to: Machine learning approach to gene essentiality prediction: a review. Brief Bioinform, 23.

Bi, S., Li, M., Liu, C., Liu, X., Cheng, J., Wang, L., Wang, J., Lv, Y., He, M., Cheng, X., Gao, Y. & Wang, C. 2022. Actin depolymerizing factor ADF7 inhibits actin bundling protein VILLIN1 to regulate root hair formation in response to osmotic stress in Arabidopsis. PLoS Genet, 18, e1010338.

Bolger, A. M., Lohse, M. & Usadel, B. 2014. Trimmomatic: a flexible trimmer for Illumina sequence data. Bioinformatics, 30, 2114–20.

Bou Daher, F., VAN Oostende, C. & Geitmann, A. 2011. Spatial and temporal expression of actin depolymerizing factors ADF7 and ADF10 during male gametophyte development in Arabidopsis thaliana. Plant Cell Physiol, 52, 1177–92.

Breiman, L., Friedman, J. H., Olshen, R., & Stone, C 1984. Classification and regression trees. Pacific Grove: Wadsworth & Brooks.

BURGOS-Rivera, B., Ruzicka, D. R., Deal, R. B., Mckinney, E. C., KING-Reid, L. & Meagher, R. B. 2008. ACTIN DEPOLYMERIZING FACTOR9 controls development and gene expression in Arabidopsis. Plant Mol Biol, 68, 619–32.

Cannon, S. B., Mitra, A., Baumgarten, A., Young, N. D. & May, G. 2004. The roles of segmental and tandem gene duplication in the evolution of large gene families in Arabidopsis thaliana. BMC Plant Biol, 4, 10.

Chen, C., Chen, H., Zhang, Y., Thomas, H. R., Frank, M. H., He, Y. & Xia, R. 2020. TBtools: An Integrative Toolkit Developed for Interactive Analyses of Big Biological Data. Mol Plant, 13, 1194–1202.

Clement, M., Ketelaar, T., Rodiuc, N., Banora, M. Y., Smertenko, A., Engler, G., Abad, P., Hussey, P. J. & De Almeida Engler, J. 2009. Actin-depolymerizing factor2-mediated actin dynamics are essential for root-knot nematode infection of Arabidopsis. Plant Cell, 21, 2963–79.

Daher, F. B. & Geitmann, A. 2012. Actin depolymerizing factors ADF7 and ADF10 play distinct roles during pollen development and pollen tube growth. Plant Signal Behav, 7, 879–81.

Dasarathy, B. V. 1991. Nearest neighbour (NN) norms: NN pattern classification techniques. IEEE Comput. Soc. Tutor, 17, 441–458.

Day, B., Henty, J. L., Porter, K. J. & Staiger, C. J. 2011. The pathogen-actin connection: a platform for defense signaling in plants. Annu Rev Phytopathol, 49, 483–506.

Dong, C. H., Xia, G. X., Hong, Y., Ramachandran, S., Kost, B. & Chua, N. H. 2001. ADF proteins are involved in the control of flowering and regulate F-actin organization, cell expansion, and organ growth in Arabidopsis. Plant Cell, 13, 1333–46.

Emms, D. M. & Kelly, S. 2015. OrthoFinder: solving fundamental biases in whole genome comparisons dramatically improves orthogroup inference accuracy. Genome Biol, 16, 157.

Emms, D. M. & Kelly, S. 2019. OrthoFinder: phylogenetic orthology inference for comparative genomics. Genome Biol, 20, 238.

Feng, Y., Liu, Q. & Xue, Q. 2006. Comparative study of rice and Arabidopsis actin-depolymerizing factors gene families. J Plant Physiol, 163, 69–79.

Hardesty, L. 2017. Explained: Neural networks. MIT News Office. Retrieved 2 *June 2022*.

He, Z., Luo, Y., Zhou, X., Zhu, T., Lan, Y. & Chen, D. 2024. scPlantDB: a comprehensive database for exploring cell types and markers of plant cell atlases. Nucleic Acids Res, 52, D1629–D1638.

Henty, J. L., Bledsoe, S. W., Khurana, P., Meagher, R. B., Day, B., Blanchoin, L. & Staiger, C. J. 2011. Arabidopsis actin depolymerizing factor4 modulates the stochastic dynamic behavior of actin filaments in the cortical array of epidermal cells. Plant Cell, 23, 3711–26.

Henty-Ridilla, J. L., Li, J., Blanchoin, L. & Staiger, C. J. 2013. Actin dynamics in the cortical array of plant cells. Curr Opin Plant Biol, 16, 678–87.

Henty-Ridilla, J. L., Li, J., Day, B. & Staiger, C. J. 2014. ACTIN DEPOLYMERIZING FACTOR4 regulates actin dynamics during innate immune signaling in Arabidopsis. Plant Cell, 26, 340–52.

Higaki, T., Sano, T. & Hasezawa, S. 2007. Actin microfilament dynamics and actin side-binding proteins in plants. Curr Opin Plant Biol, 10, 549–56.

Ho, T. K. 1995. Random Decision Forests. Proceedings of the 3rd International Conference on Document Analysis and Recognition. Montreal, Qc, pp. 278–282.

Holland, P. W., Marletaz, F., Maeso, I., Dunwell, T. L. & Paps, J. 2017. New genes from old: asymmetric divergence of gene duplicates and the evolution of development. Philos Trans R Soc Lond B Biol Sci, 372.

Horton, P., Park, K. J., Obayashi, T., Fujita, N., Harada, H., Adams-Collier, C. J. & Nakai, K. 2007. WoLF PSORT: protein localization predictor. Nucleic Acids Res, 35, W585–7.

Huang, J., Sun, W., Ren, J., Yang, R., Fan, J., Li, Y., Wang, X., Joseph, S., Deng, W. & Zhai, L. 2020. Genome-Wide Identification and Characterization of Actin-Depolymerizing Factor (ADF) Family Genes and Expression Analysis of Responses to Various Stresses in Zea Mays L. Int J Mol Sci, 21.

Huang, S., Qu, X. & Zhang, R. 2015. Plant villins: versatile actin regulatory proteins. J Integr Plant Biol, 57, 40–9.

Hussey, P. J., Ketelaar, T. & Deeks, M. J. 2006. Control of the actin cytoskeleton in plant cell growth. Annu Rev Plant Biol, 57, 109–25.

Inada, N. 2018. Correction to: Plant actin depolymerizing factor: actin microfilament disassembly and more. J Plant Res, 131, 567.

Inada, N., Higaki, T. & Hasezawa, S. 2016. Nuclear Function of Subclass I Actin-Depolymerizing Factor Contributes to Susceptibility in Arabidopsis to an Adapted Powdery Mildew Fungus. Plant Physiol, 170, 1420–34.

Inada, N., Takahashi, N. & Umeda, M. 2021. Arabidopsis thaliana subclass I ACTIN DEPOLYMERIZING FACTORs and vegetative ACTIN2/8 are novel regulators of endoreplication. J Plant Res, 134, 1291–1300.

Jiang, Y., Wang, J., Xie, Y., Chen, N. & Huang, S. 2017. ADF10 shapes the overall organization of apical actin filaments by promoting their turnover and ordering in pollen tubes. J Cell Sci, 130, 3988–4001.

Khatun, K., Robin, A. H., Park, J. I., Kim, C. K., Lim, K. B., Kim, M. B., Lee, D. J., Nou, I. S. & Chung, M. Y. 2016. Genome-Wide Identification, Characterization and Expression Profiling of ADF Family Genes in Solanum lycopersicum L. Genes (Basel), 7.

Klok, E. J., Wilson, I. W., Wilson, D., Chapman, S. C., Ewing, R. M., Somerville, S. C., Peacock, W. J., Dolferus, R. & Dennis, E. S. 2002. Expression profile analysis of the low-oxygen response in Arabidopsis root cultures. Plant Cell, 14, 2481–94.

Le, N. Q. K., Yapp, E. K. Y., Nagasundaram, N. & Yeh, H. Y. 2019. Classifying Promoters by Interpreting the Hidden Information of DNA Sequences via Deep Learning and Combination of Continuous FastText N-Grams. Front Bioeng Biotechnol, 7, 305.

Lescot, M., Dehais, P., Thijs, G., Marchal, K., Moreau, Y., Van De Peer, Y., Rouze, P. & Rombauts, S. 2002. Plantcare, a database of plant cis-acting regulatory elements and a portal to tools for in silico analysis of promoter sequences. Nucleic Acids Res, 30, 325–7.

Li, P., Lu, Y. J., Chen, H. & Day, B. 2020. The Lifecycle of the Plant Immune System. CRC Crit Rev Plant Sci, 39, 72–100.

Lin, H., Liang, Z. Y., Tang, H. & Chen, W. 2019. Identifying Sigma70 Promoters with Novel Pseudo Nucleotide Composition. IEEE/ACM Trans Comput Biol Bioinform, 16, 1316–1321.

Macrobbie, E. A. C. & Kurup, S. 2007. Signalling mechanisms in the regulation of vacuolar ion release in guard cells. New Phytol, 175, 630–640.

Matsumoto, T., Higaki, T., Takatsuka, H., Kutsuna, N., Ogata, Y., Hasezawa, S., Umeda, M. & Inada, N. 2023. Arabidopsis thaliana Subclass I ACTIN DEPOLYMERIZING FACTORs Regulate Nuclear Organization and Gene Expression. Plant Cell Physiol.

Mccurdy, D. W., Kovar, D. R. & Staiger, C. J. 2001. Actin and actin-binding proteins in higher plants. Protoplasma, 215, 89–104.

Meinke, D. W., Cherry, J. M., Dean, C., Rounsley, S. D. & Koornneef, M. 1998. Arabidopsis thaliana: a model plant for genome analysis. Science, 282, 662, 679–82.

Miklis, M., Consonni, C., Bhat, R. A., Lipka, V., Schulze-Lefert, P. & Panstruga, R. 2007. Barley MLO modulates actin-dependent and actin-independent antifungal defense pathways at the cell periphery. Plant Physiol, 144, 1132–43.

Nan, Q., Qian, D., Niu, Y., He, Y., Tong, S., Niu, Z., Ma, J., Yang, Y., An, L., Wan, D. & Xiang, Y. 2017. Plant Actin-Depolymerizing Factors Possess Opposing Biochemical Properties Arising from Key Amino Acid Changes throughout Evolution. Plant Cell, 29, 395–408.

Patro, R., Duggal, G., Love, M. I., Irizarry, R. A. & Kingsford, C. 2017. Salmon provides fast and bias-aware quantification of transcript expression. Nat Methods, 14, 417–419.

Pavlov, D., Muhlrad, A., Cooper, J., Wear, M. & Reisler, E. 2007. Actin filament severing by cofilin. J Mol Biol, 365, 1350–8.

Peng, S. Q. & Huang, D. F. 2006. [Expression of an Arabidopsis actin-depolymerizing factor 4 gene (AtADF4) in tobacco causes morphological change of plants]. Zhi Wu Sheng Li Yu Fen Zi Sheng Wu Xue Xue Bao, 32, 52–6.

Pollard, T. D. & Cooper, J. A. 2009. Actin, a central player in cell shape and movement. Science, 326, 1208–12.

Porter, K., Shimono, M., Tian, M. & Day, B. 2012. Arabidopsis Actin-Depolymerizing Factor-4 links pathogen perception, defense activation and transcription to cytoskeletal dynamics. PLoS Pathog, 8, e1003006.

Qian, D., Zhang, Z., He, J., Zhang, P., Ou, X., Li, T., Niu, L., Nan, Q., Niu, Y., He, W., An, L., Jiang, K. & Xiang, Y. 2019. Arabidopsis ADF5 promotes stomatal closure by regulating actin cytoskeleton remodeling in response to ABA and drought stress. J Exp Bot, 70, 435–446.

ROY-Zokan, E. M., Dyer, K. A. & Meagher, R. B. 2015. Phylogenetic Patterns of Codon Evolution in the ACTIN-DEPOLYMERIZING FACTOR/COFILIN (ADF/CFL) Gene Family. PLoS One, 10, e0145917.

Ruzicka, D. R., Kandasamy, M. K., Mckinney, E. C., BURGOS-Rivera, B. & Meagher, R. B. 2007. The ancient subclasses of Arabidopsis Actin Depolymerizing Factor genes exhibit novel and differential expression. Plant J, 52, 460–72.

Staiger, C. J. 2000. Signaling to the Actin Cytoskeleton in Plants. Annu Rev Plant Physiol Plant Mol Biol, 51, 257–288.

Staiger, C. J. & Blanchoin, L. 2006. Actin dynamics: old friends with new stories. Curr Opin Plant Biol, 9, 554–62.

Stamatakis, A. 2014. RAxML version 8: a tool for phylogenetic analysis and post-analysis of large phylogenies. Bioinformatics, 30, 1312–3.

Sun, T., Li, S. & Ren, H. 2013. Profilin as a regulator of the membrane-actin cytoskeleton interface in plant cells. Front Plant Sci, 4, 512.

Szymanski, D. B. & Cosgrove, D. J. 2009. Dynamic coordination of cytoskeletal and cell wall systems during plant cell morphogenesis. Curr Biol, 19, R800–11.

Tholl, S., Moreau, F., Hoffmann, C., Arumugam, K., Dieterle, M., Moes, D., Neumann, K., Steinmetz, A. & Thomas, C. 2011. Arabidopsis actin-depolymerizing factors (ADFs) 1 and 9 display antagonist activities. FEBS Lett, 585, 1821–7.

Tian, M., Chaudhry, F., Ruzicka, D. R., Meagher, R. B., Staiger, C. J. & Day, B. 2009. Arabidopsis actin-depolymerizing factor AtADF4 mediates defense signal transduction triggered by the Pseudomonas syringae effector AvrPphB. Plant Physiol, 150, 815–24.

VAN Gisbergen, P. A. & Bezanilla, M. 2013. Plant formins: membrane anchors for actin polymerization. Trends Cell Biol, 23, 227–33.

Vilo, J., Brazma, A., Jonassen, I., Robinson, A. & Ukkonen, E. 2000. Mining for putative regulatory elements in the yeast genome using gene expression data. Proc Int Conf Intell Syst Mol Biol, 8, 384–94.

Wang, L., Cheng, J., Bi, S., Wang, J., Cheng, X., Liu, S., Gao, Y., Lan, Q., Shi, X., Wang, Y., Zhao, X., Qi, X., Xu, S. & Wang, C. 2023. Actin Depolymerization Factor ADF1 Regulated by MYB30 Plays an Important Role in Plant Thermal Adaptation. Int J Mol Sci, 24.

Wang, Y., Tang, H., Debarry, J. D., Tan, X., Li, J., Wang, X., Lee, T. H., Jin, H., Marler, B., Guo, H., Kissinger, J. C. & Paterson, A. H. 2012. MCScanX: a toolkit for detection and evolutionary analysis of gene synteny and collinearity. Nucleic Acids Res, 40, e49.

Wigge, P. A. & Weigel, D. 2001. Arabidopsis genome: life without notch. Curr Biol, 11, R112–4.

Wu, L. & Yang, H. Q. 2010. CRYPTOCHROME 1 is implicated in promoting R protein-mediated plant resistance to Pseudomonas syringae in Arabidopsis. Mol Plant, 3, 539–48.

Zhang, P., Qian, D., Luo, C., Niu, Y., Li, T., Li, C., Xiang, Y., Wang, X. & Niu, Y. 2021. Arabidopsis ADF5 Acts as a Downstream Target Gene of CBFs in Response to Low-Temperature Stress. Front Cell Dev Biol, 9, 635533.

Zhao, S., Jiang, Y., Zhao, Y., Huang, S., Yuan, M., Zhao, Y. & Guo, Y. 2016. CASEIN KINASE1-LIKE PROTEIN2 Regulates Actin Filament Stability and Stomatal Closure via Phosphorylation of Actin Depolymerizing Factor. Plant Cell, 28, 1422–39.

Zhao, Y., Zhao, S., Mao, T., Qu, X., Cao, W., Zhang, L., Zhang, W., He, L., Li, S., Ren, S., Zhao, J., Zhu, G., Huang, S., Ye, K., Yuan, M. & Guo, Y. 2011. The plant-specific actin binding protein SCAB1 stabilizes actin filaments and regulates stomatal movement in Arabidopsis. Plant Cell, 23, 2314–30.

Zheng, Y., Xie, Y., Jiang, Y., Qu, X. & Huang, S. 2013. Arabidopsis actin-depolymerizing factor7 severs actin filaments and regulates actin cable turnover to promote normal pollen tube growth. Plant Cell, 25, 3405–23.

Zhu, J., Nan, Q., Qin, T., Qian, D., Mao, T., Yuan, S., Wu, X., Niu, Y., Bai, Q., An, L. & Xiang, Y. 2017. Higher-Ordered Actin Structures Remodeled by Arabidopsis ACTIN-DEPOLYMERIZING FACTOR5 Are Important for Pollen Germination and Pollen Tube Growth. Mol Plant, 10, 1065–1081.

